# Antiviral treatment of SARS-CoV-2-infected hamsters reveals a weak effect of favipiravir and a complete lack of effect for hydroxychloroquine

**DOI:** 10.1101/2020.06.19.159053

**Authors:** Suzanne JF Kaptein, Sofie Jacobs, Lana Langendries, Laura Seldeslachts, Sebastiaan ter Horst, Laurens Liesenborghs, Bart Hens, Valentijn Vergote, Elisabeth Heylen, Elke Maas, Carolien De Keyzer, Lindsey Bervoets, Jasper Rymenants, Tina Van Buyten, Hendrik Jan Thibaut, Kai Dallmeier, Robbert Boudewijns, Jens Wouters, Patrick Augustijns, Nick Verougstraete, Christopher Cawthorne, Birgit Weynand, Pieter Annaert, Isabel Spriet, Greetje Vande Velde, Johan Neyts, Joana Rocha-Pereira, Leen Delang

## Abstract

SARS-CoV-2 rapidly spread around the globe after its emergence in Wuhan in December 2019. With no specific therapeutic and prophylactic options available, the virus was able to infect millions of people. To date, close to half a million patients succumbed to the viral disease, COVID-19. The high need for treatment options, together with the lack of small animal models of infection has led to clinical trials with repurposed drugs before any preclinical *in vivo* evidence attesting their efficacy was available. We used Syrian hamsters to establish a model to evaluate antiviral activity of small molecules in both an infection and a transmission setting. Upon intranasal infection, the animals developed high titers of SARS-CoV-2 in the lungs and pathology similar to that observed in mild COVID-19 patients. Treatment of SARS-CoV-2-infected hamsters with favipiravir or hydroxychloroquine (with and without azithromycin) resulted in respectively a mild or no reduction in viral RNA and infectious virus. Micro-CT scan analysis of the lungs showed no improvement compared to non-treated animals, which was confirmed by histopathology. In addition, both compounds did not prevent virus transmission through direct contact and thus failed as prophylactic treatments. By modelling the PK profile of hydroxychloroquine based on the trough plasma concentrations, we show that the total lung exposure to the drug was not the limiting factor. In conclusion, we here characterized a hamster infection and transmission model to be a robust model for studying *in vivo* efficacy of antiviral compounds. The information acquired using hydroxychloroquine and favipiravir in this model is of critical value to those designing (current and) future clinical trials. At this point, the data here presented on hydroxychloroquine either alone or combined with azithromycin (together with previously reported *in vivo* data in macaques and ferrets) provide no scientific basis for further use of the drug in humans.

## Introduction

The severe acute respiratory syndrome coronavirus 2 (SARS-CoV-2) first emerged in Wuhan, China in December 2019^1^. From there, the virus rapidly spread around the globe, infecting more than 8 million people so far (June 18) [https://covid19.who.int/]. SARS-CoV-2 is the causative agent of coronavirus disease 2019 (COVID-19). Common clinical manifestations of COVID-19 are fever, dry cough, paired in a minority of patients with difficult breathing, muscle and/or joint pain, headache/dizziness, decreased sense of taste and smell, diarrhea, and nausea^2^. A small subset of patients will develop to acute respiratory distress syndrome (ARDS), characterized by difficult breathing and low blood oxygen levels, which may directly result into respiratory failure^2^. In addition, an overreaction of the host’s immune and inflammatory responses can result in a vast release of cytokines (‘cytokine storm’), inducing sepsis and multi-organ damage, which may lead to organ failure^3^. To date, more than 440,000 patients worldwide succumbed to COVID-19. Hence, in response to the ongoing pandemic there is a desperate need for therapeutic and prophylactic options.

At present, no specific antiviral drugs have been developed and approved to treat infections with human coronaviruses. Nonetheless, antiviral drugs could fulfill an important role in the treatment of COVID-19 patients. Slowing down the replication of SARS-CoV-2 by antiviral treatment could be beneficial and prevent or alleviate symptoms. In addition, antiviral drugs could be used as prophylaxis to protect health care workers and high-risk groups. However, a specific, highly potent antiviral drug for SARS-CoV-2 will take years to develop and evaluate in clinical studies. Therefore, the main focus for COVID-19 treatment on the short term is on the repurposing of drugs that have been approved for other diseases^4^. Repurposed drugs can however not be expected to be highly potent inhibitors of SARS-CoV-2, since these were not developed and optimized specifically against this virus. In cell culture, several repurposed drugs inhibit SARS-CoV-2 replication^5,6^. Although preclinical *in vivo* evidence evaluating the efficacy of some of these repurposed drugs for COVID-19 treatment is lacking, clinical trials have already been conducted or are currently ongoing. Two such drugs are hydroxychloroquine and favipiravir.

Hydroxychloroquine (HCQ) is an anti-malaria drug that has been widely used to treat patients with malaria, rheumatoid arthritis and systemic lupus erythematosus. This drug is also able to inhibit a broad range of viruses from different virus families in cell culture, including coronaviruses (SARS-CoV-1, MERS-CoV)^7,8^. Favipiravir is a broad-spectrum antiviral drug that has been approved in Japan since 2014 to treat pandemic influenza virus infections^9^. Both drugs have shown antiviral efficacy against SARS-CoV-2 in Vero E6 cells^10^, albeit modest for favipiravir^10–12^. Enzymatic assays with the SARS-CoV-2 RNA-dependent RNA polymerase demonstrated that favipiravir acts as a nucleotide analog via a combination of chain termination, slowed viral RNA synthesis and lethal mutagenesis^12^. However, proof of *in vivo* efficacy in animal models is still lacking for both drugs. Nevertheless, clinical trials were initiated early on in the pandemic to assess the efficacy of HCQ and favipiravir to treat COVID-19 patients. For HCQ, these trials were small anecdotal studies or inconclusive small randomized trials^13^ and thus did not lead to conclusive results. Despite the lack of clear evidence, HCQ is currently being widely used for the treatment of COVID-19, often in combination with a second-generation macrolide such as azithromycin. Results from animal models and rigorous randomized controlled trials are thus required to clarify the efficacy of HCQ and favipiravir in the treatment of COVID-19 patients.

Infection models in small animals are crucial for the evaluation and development of antiviral drugs. Although rhesus and cynomolgus macaques seem to be relevant models for studying the early stages of COVID-19 infection in humans^14^, preclinical models using smaller animals are essential to ensure efficient and ethical allocation of resources towards designing (relevant) preclinical and clinical efficacy studies. Syrian hamsters are permissive to SARS-CoV-2 and develop mild lung disease similar to the disease observed in early-stage COVID-19 patients^15,16^. Nevertheless, evidence of antiviral efficacy of repurposed drugs in small animal models is lacking to date. In this work, we characterized Syrian hamsters as a model for the evaluation of antiviral drugs in therapeutic and prophylactic settings against SARS-CoV-2. We then used this model to evaluate the antiviral efficacy of HCQ and favipiravir against SARS-CoV-2 in infected hamsters and in a transmission setting.

## Material and Methods

### SARS-CoV-2

The SARS-CoV-2 strain used in this study, BetaCov/Belgium/GHB-03021/2020 (EPI ISL 407976|2020-02-03), was recovered from a nasopharyngeal swab taken from a RT-qPCR-confirmed asymptomatic patient who returned from Wuhan, China in the beginning of February 2020^17^. A close relation with the prototypic Wuhan-Hu-1 2019-nCoV (GenBank accession number MN908947.3) strain was confirmed by phylogenetic analysis. Infectious virus was isolated by serial passaging on HuH7 and Vero E6 cells^16^; passage 6 virus was used for the studies described here. The titer of the virus stock was determined by end-point dilution on Vero E6 cells by the Reed and Muench method. Live virus-related work was conducted in the high-containment A3 and BSL3^+^ facilities of the KU Leuven Rega Institute (3CAPS) under licenses AMV 30112018 SBB 219 2018 0892 and AMV 23102017 SBB 219 20170589 according to institutional guidelines.

### Cells

Vero E6 cells (African green monkey kidney, ATCC CRL-1586) were cultured in minimal essential medium (Gibco) supplemented with 10% fetal bovine serum (Integro), 1% L-glutamine (Gibco) and 1% bicarbonate (Gibco). End-point titrations were performed with medium containing 2% fetal bovine serum instead of 10%.

### Compounds

Favipiravir was purchased from BOC Sciences (USA). Hydroxychloroquine sulphate was acquired from Acros Organics. For *in vivo* treatment, a 30 mg/mL favipiravir suspension was prepared in 0.4% carboxymethylcellulose and a 20 mg/mL hydroxychloroquine sulphate solution in 10% DMSO, 18% Cremophor, and 72% water. Azithromycin was provided by the hospital pharmacy of the University Hospitals Leuven (Belgium) as a 40 mg/ml oral solution (Zitromax^®^) which was diluted to 5 mg/mL with an aqueous medium consisting of 0.6% xanthan gum as viscosity enhancer.

### SARS-CoV-2 infection model in hamsters

The hamster infection model of SARS-CoV-2 has been described before^16^. In brief, wild-type Syrian hamsters (*Mesocricetus auratus*) were purchased from Janvier Laboratories and were housed per two in ventilated isolator cages (IsoCage N Biocontainment System, Tecniplast) with *ad libitum* access to food and water and cage enrichment (wood block). Housing conditions and experimental procedures were approved by the ethical committee of animal experimentation of KU Leuven (license P065-2020).

Female hamsters of 6-10 weeks old were anesthetized with ketamine/xylazine/atropine and inoculated intranasally with 50 μL containing 2×10^6^ TCID_50_. Drug treatment was initiated 1h before infection. Favipiravir was administered twice daily by oral gavage, starting with a loading dose of 600 mg/kg/day on the first day. On consecutive days, 300 mg/kg/day favipiravir was administered until the day of sacrifice. Hydroxychloroquine sulphate (50 mg/kg) was administered once daily by intraperitoneal (ip) injection for 4 days. Azithromycin (10 mg/kg) was administered once daily by oral gavage using a 5 mg/ml dilution of Zitromax^®^. Hamsters were daily monitored for appearance, behavior and weight. At day 4 post infection (pi), hamsters were euthanized by ip injection of 500 μL Dolethal (200mg/mL sodium pentobarbital, Vétoquinol SA). Tissues [lungs, small intestine (ileum)] and stool were collected, and viral RNA and infectious virus were quantified by RT-qPCR and end-point virus titration, respectively. Blood samples were collected at day 4 pi for PK analysis of HCQ.

### SARS-CoV-2 transmission model in hamsters

The hamster transmission model of SARS-CoV-2 via direct contact has been described previously^15,18^. Briefly, index hamsters (6-10 weeks old) were infected as described above. At the day of exposure, sentinel hamsters were co-housed with index hamsters that had been intranasally inoculated with SARS-CoV-2 one day earlier. Index and sentinel hamsters were sacrificed at day 4 pi (post-exposure in the case of the sentinels) and the viral load in lung, ileum and stool was determined, as described above. For prophylactic testing of drugs, sentinel hamsters were treated daily for 5 consecutive days with either hydroxychloroquine or favipiravir, starting 1 day prior to exposure to the index hamster.

To study the contribution of the fecal-oral route to the overall transmission of SARS-CoV-2, index hamsters were inoculated as described earlier. On day 1 or 3 pi, the index hamsters were sacrificed after which sentinel hamsters were placed in the dirty cages of the index hamsters. Food grids and water bottles were replaced by clean ones to minimize virus transmission via food or water. At day 4 post exposure, the sentinels were sacrificed. Tissues (lung, ileum and stool) were collected from index and sentinel hamsters and processed for detection of viral RNA and infectious virus.

### PK analysis of hydroxychloroquine and metabolite in plasma

Hydroxychloroquine (HCQ) and its active metabolite desethylhydroxychloroquine (DHCQ) were quantified in EDTA-plasma samples. A total of (i) 50 μL sample and (ii) 10 μL of internal standard (IS) solution (hydroxychloroquine-d4 1500 ng/mL in water) were added to a tube and mixed. After addition of 50 μL 5% perchloric acid, samples were shaken for 5 min and centrifuged for 5 min at 16,162 g. Five μL of the supernatant was injected onto the HPLC-column.

HPLC analysis was performed using a Shimadzu Prominence system (Shimadzu, Kioto, Japan) equipped with a Kinetex C18 column (100mm length x 2.1mm i.d., 2.6 μm particle size) (Phenomenex, Torrance, CA, USA) at 50°C. A 6 min gradient of mobile phase A (0.1% formic acid (FA) in water) and B (0.1% FA in acetonitrile) with a flow rate of 0.4 mL/min was used for elution of the compounds. The mass spectrometer (MS) was a Triple Quad 5500 (Sciex, Framingham, MA, USA) with an electrospray ionization source (ESI) in positive ion mode, using multiple reaction monitoring (MRM). The monitored transitions were 336.8 to 248.0 m/z, 307.8 to 130.0 m/z and 340.8 to 252.0 m/z for HCQ, DHCQ, and HCQ-d4, respectively. The used collision energy for all the transitions was 30 V. Calibration curves for both HCQ (linear 1/x weighting) and DHCQ (quadratic 1/x^2^ weighting) were between 10 and 2250 ng/mL. Between-run imprecision over all QC levels (10, 25, 400, 2000 ng/mL) ranged from 2.84 to 11.4% for HCQ and from 5.19 to 10.2% for DHCQ.

### Calculation of hydroxychloroquine concentration in the lung cytosol

Starting from the measured total trough plasma concentrations measured at sacrifice after 4 or 5 days of HCQ treatment, total lung cytosolic concentrations of HCQ were calculated. First, the mean trough total plasma concentration of HCQ was used as a starting point to estimate the whole blood concentrations considering a blood to plasma ratio of 7.2, as reported by Tett and co-workers^19^ and as mentioned in the SmPC of Plaquenil^®^ (Sanofi, Paris, France) (**Equation 1**).

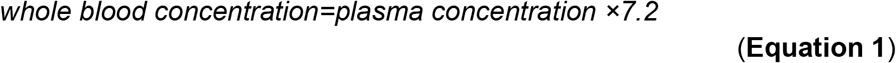

Relying on the experimental Kp (tissue versus whole blood partition coefficient) values in rats, the total lung tissue concentrations of HCQ was determined. Based on the partition values as reported by Wei et al.^20^, a lung Kp value of 50 was applied to estimate the total lung concentration (**Equation 2**).

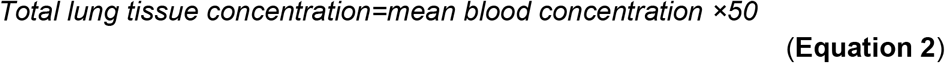

Subsequently, as the HCQ efficacy target is intracellular, the cytosolic / total HCQ concentration ratio was estimated, based on (i) relative lysosomal lung tissue volume, as well as the contributions of interstitial and intracellular volumes to total lung volume and (ii) the pH partition theory applying a pKa value of HCQ of 9.67. Based on these calculations (data not shown), lung cytosolic HCQ concentrations are corresponding to 6% of the total lung tissue concentration (**Equation 3**).

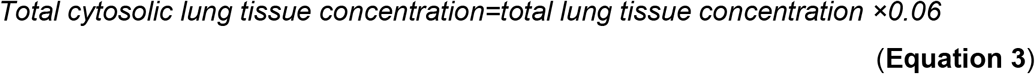

The calculated total cytosolic lung concentration was compared with EC_50_ concentrations previously reported in literature, ranging from 0.72 μM to 17.3 μM^21–23^.

### SARS-CoV-2 RT-qPCR

Hamster tissues were collected after sacrifice and were homogenized using bead disruption (Precellys) in 350 μL RLT buffer (RNeasy Mini kit, Qiagen) and centrifuged (10,000 rpm, 5 min) to pellet the cell debris. RNA was extracted according to the manufacturer’s instructions. To extract RNA from serum, the NucleoSpin kit (Macherey-Nagel) was used. Of 50 μL eluate, 4 μL was used as a template in RT-qPCR reactions. RT-qPCR was performed on a LightCycler96 platform (Roche) using the iTaq Universal Probes One-Step RT-qPCR kit (BioRad) with N2 primers and probes targeting the nucleocapsid^16^. Standards of SARS-CoV-2 cDNA (IDT) were used to express viral genome copies per mg tissue or per mL serum.

### End-point virus titrations

Lung tissues were homogenized using bead disruption (Precellys) in 350 μL minimal essential medium and centrifuged (10,000 rpm, 5min, 4°C) to pellet the cell debris. To quantify infectious SARS-CoV-2 particles, endpoint titrations were performed on confluent Vero E6 cells in 96-well plates. Viral titers were calculated by the Reed and Muench method using the Lindenbach calculator^24^ and were expressed as 50% tissue culture infectious dose (TCID_50_) per mg tissue.

### Histology

For histological examination, the lungs were fixed overnight in 4% formaldehyde and embedded in paraffin. Tissue sections (5 μm) were analyzed after staining with hematoxylin and eosin and scored blindly for lung damage by an expert pathologist. The scored parameters, to which a cumulative score of 1 to 3 was attributed, were the following: congestion, intral-alveolar hemorrhagic, apoptotic bodies in bronchus wall, necrotizing bronchiolitis, perivascular edema, bronchopneumonia, perivascular inflammation, peribronchial inflammation and vasculitis.

### Micro-computed tomography (CT) and image analysis

Micro-CT data of hamster lungs were acquired *in vivo* using dedicated small animal micro-CT scanners, either using the X-cube (Molecubes, Ghent, Belgium) or the Skyscan 1278 (Bruker Belgium, Kontich, Belgium). In brief, hamsters were anaesthetized using isoflurane (2-3% in oxygen) and installed in prone position into the X-cube scanner using a dedicated imaging bed. A scout view was acquired and the lung was selected for a non-gated, helical CT acquisition using the High-Resolution CT protocol, with the following parameters: 50 kVp, 960 exposures, 32 ms/projection, 350 μA tube current, rotation time 120 s. Data were reconstructed with 100 μm isotropic voxel size using a regularized statistical (iterative) image reconstruction algorithm^25^. On the SkyScan1278, hamsters were scanned in supine position under isoflurane anesthesia and the following scan parameters were used: 55 kVp X-ray source voltage and 500 μA current combined with a composite X-ray filter of 1 mm aluminium, 80 ms exposure time per projection, acquiring 4 projections per step with 0.7° increments over a total angle of 220°, and 10 cm field of view covering the whole body producing expiratory weighted 3D data sets with 50 μm isotropic reconstructed voxel size^26^. Each scan took approximately 3 minutes.

Visualization and quantification of reconstructed micro-CT data were performed with DataViewer and CTan software (Bruker Belgium). As primary outcome measure, a semi-quantitative scoring of micro-CT data was performed as previously described^25^. Visual observations were blindly scored (from 0 – 2 depending on severity, both for parenchymal and airway disease) on 5 different, predefined transversal tomographic sections throughout the entire lung image for both lung and airway disease by two independent observers and averaged. Scores for the 5 sections were summed up to obtain a score from 0 to 10 reflecting severity of lung and airway abnormalities compared to scans of healthy, WT control hamsters. As secondary measures, imaging-derived biomarkers (non-aerated lung volume, aerated lung volume, total lung volume and respective densities within these volumes) were quantified as previously^16,26,27^ or a manually delineated volume of interest covering the lung, avoiding the heart and main blood vessels. The threshold used to distinguish aerated from non-aerated lung volume was manually defined and kept constant for all data sets^26,27^.

### Statistics

GraphPad Prism (GraphPad Software, Inc.). was used to perform statistical analysis. Statistical significance was determined using the non-parametric Mann Whitney U-test. P values of ≤0.05 were considered significant.

## Results

### Characterization of hamster model for antiviral drug evaluation

We further characterized SARS-CoV-2 infection and readouts of disease in hamsters to be able to use this model for the evaluation and development of antiviral drugs. To investigate SARS-CoV-2 replication and shedding, the lung, ileum and stool of infected hamsters were harvested at different time points post-infection (pi) for viral RNA quantification by RT-qPCR. Infectious virus titers were additionally determined in lung samples. SARS-CoV-2 efficiently replicates in the lungs of the hamsters, with viral RNA being detected in the lungs from day 1 pi and reaching a maximum level of ~7 log_10_ RNA copies/mg tissue at 4 days pi (Fig 1A). A similar kinetic profile was found in the ileum and stool samples, albeit at lower levels of 2-3 log_10_ RNA copies/mg of tissue. Titrations of homogenized lung tissue contained infectious particles from 1 day pi and reached levels of ~5 log_10_ TCID_50_/mg tissue from day 2 pi onwards (Fig 1B), which is in line with the viral RNA levels. Infected animals displayed a slight weight loss of about 5% by day 2 pi, which was completely resolved by day 4 pi (Fig 1C). No other signs of disease or distress were observed in the hamsters at any time point pi.

**Figure 1.**
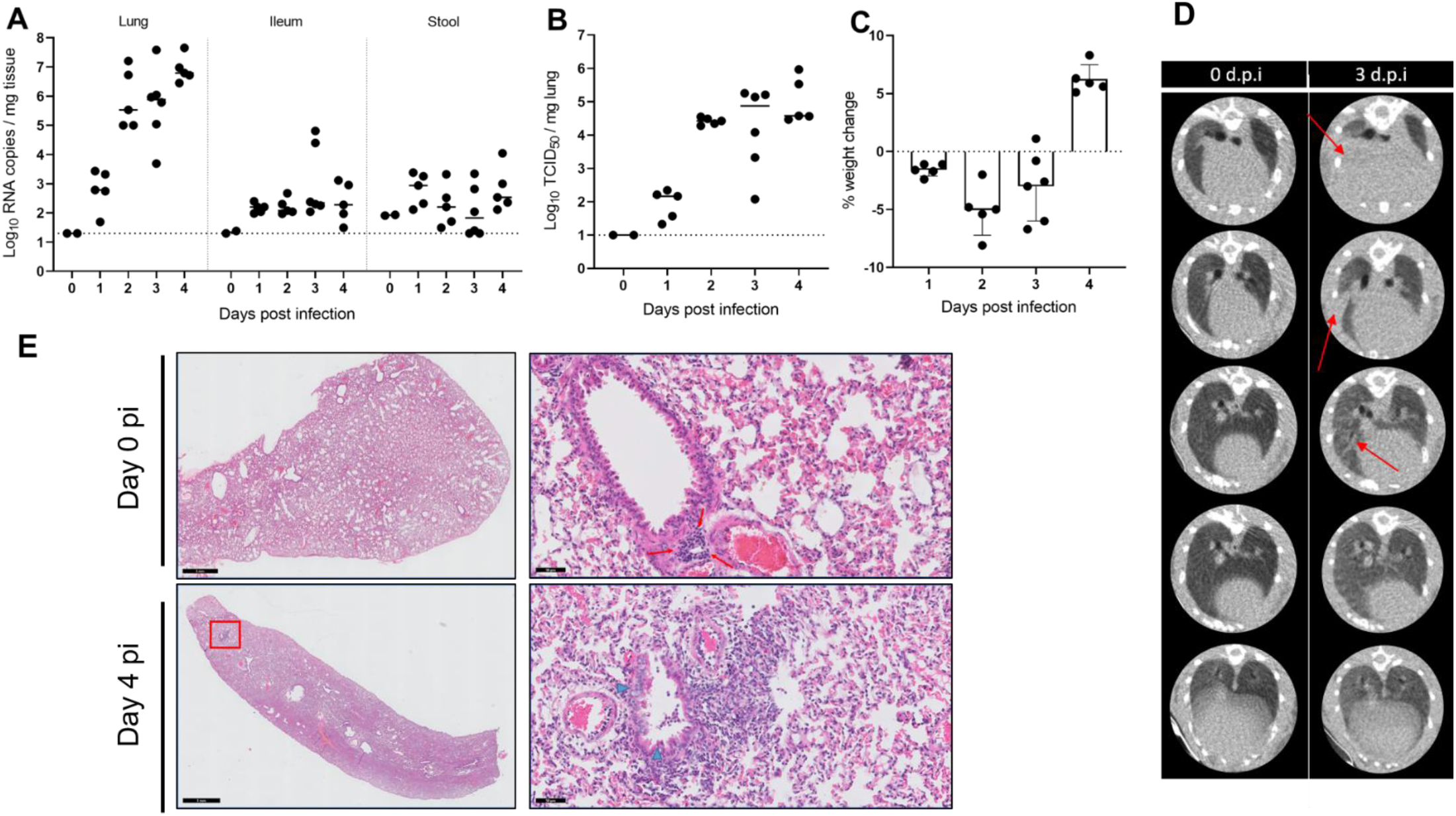
Kinetics of SARS-CoV-2 replication and lung disease in hamsters. **(A)** Viral RNA levels in the lungs, ileum and stool of infected Syrian hamsters. At the indicated time intervals pi, viral RNA levels were quantified by RT-qPCR. **(B)** Infectious viral load in the lung expressed as TCID_50_ per mg of lung tissue obtained at day 4 pi. **(C)** Weight change as compared to the weight at d0 in percentage at the indicated time intervals pi. **(A-C)** The data shown are medians plus the individual hamsters represented as separate data points. **(D)** Representative transversal lung μCT-images on SARS-CoV-2 infected hamsters at baseline (0 d.p.i) and 3 d.p.i. Red arrows indicate infiltration by consolidation of lung parenchyma. **(E)** Representative H&E images of lungs of SARS-CoV-2-infected hamsters at day 0 and day 4 pi. Red arrows point at a lymphoid follicle. The blue arrowhead indicates apoptotic cells in the bronchial epithelium.

Alike to what is currently done in clinical practice, we evaluated the development of lung disease in a non-invasive way by micro-computed tomography (micro-CT) scanning the infected animals under isoflurane gas anesthesia^28^. Dense lung infiltrations and bronchial dilation were simultaneously present from day 3 pi onwards, becoming more pronounced at day 4 pi. Longitudinal follow-up of radiological pathology showed signs of multifocal pulmonary infiltrates and lung consolidation on day 3 pi (Fig 1D). Analysis by H&E staining of lungs of infected hamsters at day 4 pi showed signs of bronchopneumonia and peribronchial inflammation, which were not present at the day of inoculation (Fig 1E).

### Evaluation of in vivo efficacy of hydroxychloroquine and favipiravir

Next, we treated hamsters with antiviral molecules for four consecutive days starting one hour before intranasal infection with SARS-CoV-2. At day 4 pi, a micro-CT scan was performed, after which the animals were sacrificed and organs were collected for quantification of viral RNA, infectious virus titers and lung histopathology (Fig 2A). Twice-daily treatment with favipiravir was done orally with a loading dose of 600 mg/kg/day at day 0 pi and 300 mg/kg/day from day 1 pi onwards. Favipiravir-treated hamsters presented a decrease of 0.9 log_10_ RNA copies/mg lung tissue, compared to untreated infected hamsters (Fig 2B); a lesser effect was observed in the ileum and stool of treated animals (Fig 2B). A modest reduction in infectious titers of 0.5 log_10_ TCID_50_/mg was observed in the lungs of favipiravir-treated animals (Fig. 2C). Treatment with favipiravir caused over 5% weight loss at day 3 and 4 pi, which is slightly more than that of the untreated animals (Fig. 2D). This could be due to the effect of administering a relatively high volume of compound per os (which was at the limit of 10 mL/kg/ day) or due to some toxicity of the molecule. Despite the very modest reduction in viral load, no obvious change (improvement or worsening) of the rather subtle radiological and histological lung pathology could be observed in favipiravir-treated hamsters (Fig. 2E-G). Quantification of micro-CT-derived biomarkers support these observations and quantify a relatively small burden of radiological lung consolidation upon infection that does not change with favipiravir treatment (Fig. 2F).

**Figure 2.**
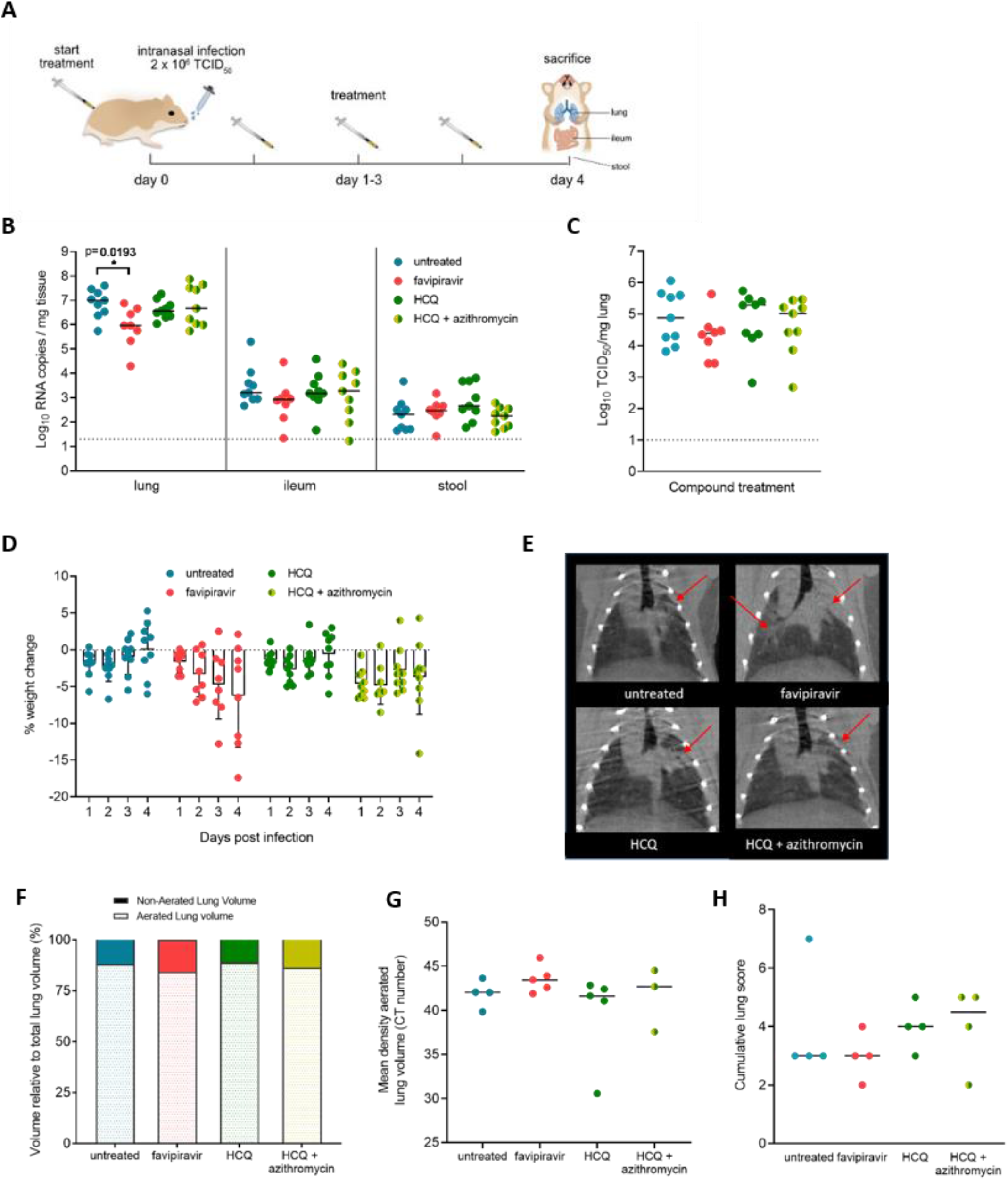
*In vivo* testing of favipiravir and hydroxychloroquine (HCQ) in the SARS-CoV-2 infection model. **(A)** Set-up of the study. **(B)** Viral RNA levels in the lungs, ileum and stool of untreated and treated (favipiravir, HCQ or HCQ + azithromycin) SARS-CoV-2 infected hamsters at day 4 pi. At the indicated time intervals pi, viral RNA levels were quantified by RT-qPCR. **(C)** Infectious viral load in the lung of untreated hamsters and hamsters receiving treatment (favipiravir, HCQ or HCQ + azithromycin) expressed as TCID_50_ per mg of lung tissue obtained at day 4 pi. **(D)** Weight change of the hamsters as compared to the weight at d0 in percentage points at the indicated time intervals pi. **(E)** Coronal lung μCT images at 4 d.p.i. of SARS-CoV-2 infected hamsters, untreated and treated with favipiravir, HCQ or HCQ + azithromycin. Red arrows point to examples of pulmonary infiltrates observed as consolidation of lung parenchyma. **(F, G)** Quantification of μCT-derived biomarkers: non-aerated lung volume (reflecting the tissue lesion volume) and aerated lung volume relative to total lung volume **(F)** and mean density of the aerated lung volume **(G)**. **(H)** Cumulative severity score from H&E staining of lungs of SARS-CoV-2 infected hamsters that were untreated (blue) or treated with favipiravir (red), HCQ (green) or HCQ + azithromycin (green-yellow).

HCQ sulphate was tested alone or in combination with azithromycin at a dose of 50 mg/kg/day (equivalent to 39 mg/kg HCQ base) administered intraperitoneally once daily. When in combination, azithromycin was given orally once daily at a dose of 10 mg/kg/day. Treatment with HCQ alone resulted in a very modest reduction of 0.3 log_10_ viral RNA copies/mg lung, and no reduction in viral RNA load in the ileum or stool compared to untreated infected hamsters (Fig 2B). When combined with azithromycin, no additional reduction of viral RNA was observed in the organs of infected animals (Fig 2B). Virus titrations of the lungs also revealed no significant reduction after treatment with HCQ alone or in combination with azithromycin (Fig 2C). The weight loss of the animals treated with HCQ follows along the lines of the untreated animals with < 5% weight loss during the whole experiment, while the combination treatment with azithromycin caused a slightly greater weight loss at day 1 and 2 pi, from which the animals could partially recover (Fig 2D). Similarly, no radiological improvement was observed between non-treated animals and animals treated with HCQ or HCQ in combination with azithromycin, which was confirmed by quantification of micro-CT-derived biomarkers of lung pathology (Fig 2E-G).

### Hydroxychloroquine and favipiravir fail to prevent SARS-CoV-2 infection in a transmission model

SARS-CoV-2 is typically transmitted through direct contact with respiratory droplets of an infected person or from touching eyes, nose or mouth after touching virus-contaminated surfaces. Transmission of SARS-CoV-2 through aerosols and direct contact has also been demonstrated in a Syrian hamster model^15,18^. We additionally explored whether SARS-CoV-2 can be transmitted via the fecal-oral route. To this end, hamsters that were intranasally inoculated with virus were sacrificed at day 1 or day 3 pi. Subsequently, sentinel hamsters were housed in the used cages of the index hamsters (food grids and water bottles were replaced by fresh ones) and sacrificed at day 4 post exposure. Although viral RNA and infectious virus could readily be detected in tissues from index hamsters (except in two stool samples), the majority of sentinel hamsters did not become infected, as shown by the absence of viral RNA and infectious virus in lung and ileum. (Supplemental Fig. 1). This indicates that the fecal-oral route only marginally contributes to the transmission SARS-Cov-2 between hamsters, thereby confirming the results of a previous study^18^. We therefore continued by focusing on transmission of the virus via direct contact only.

Using the transmission model, we investigated the prophylactic potential of HCQ and favipiravir against SARS-CoV-2. Sentinel hamsters received a daily dosage for 5 consecutive days with either HCQ or favipiravir, starting 24 hours prior to exposure. Each individual sentinel hamster was co-housed with an index hamster that had been intranasally inoculated with SARS-CoV-2 the day before (Fig 3A). Index hamsters were sacrificed 4 days pi and sentinels 4 days post exposure, after which the viral loads in lung, ileum and stool were determined. Index hamsters had ~7 log_10_ viral RNA copies/mg in the lungs, whereas untreated sentinel hamsters had ~4 log_10_ viral RNA copies/mg in the lungs (Fig 3B). Even though the variability between individual hamsters in the sentinel groups was more pronounced than in the index groups, no reduction in viral RNA was observed in either favipiravir- or HCQ-treated sentinel hamsters. Also in ileum and stool, the viral RNA levels were not reduced by compound treatment. The infectious viral loads in the lungs were also not reduced by treatment with either compound (Fig 3C), which is in line with the viral RNA data. In contrast to index hamsters, sentinel hamsters did not lose weight, but gained around 8% of body weight by day 4 pi. Sentinels that received HCQ or favipiravir treatment gained less body weight than the untreated sentinels (5% and 2%, respectively) (Fig. 3D). Pathology scores derived from micro-CT scans of hamsters revealed multifocal pulmonary infiltrates and consolidations in some but not in all hamsters (Fig. 3E, 3F). Also, micro-CT-derived biomarkers showed no difference in lung pathology between untreated and treated sentinel hamsters (Fig. 3G), further confirming that hydroxychloroquine and favipiravir failed to prevent SARS-CoV-2 infection in a transmission model.

**Figure 3.**
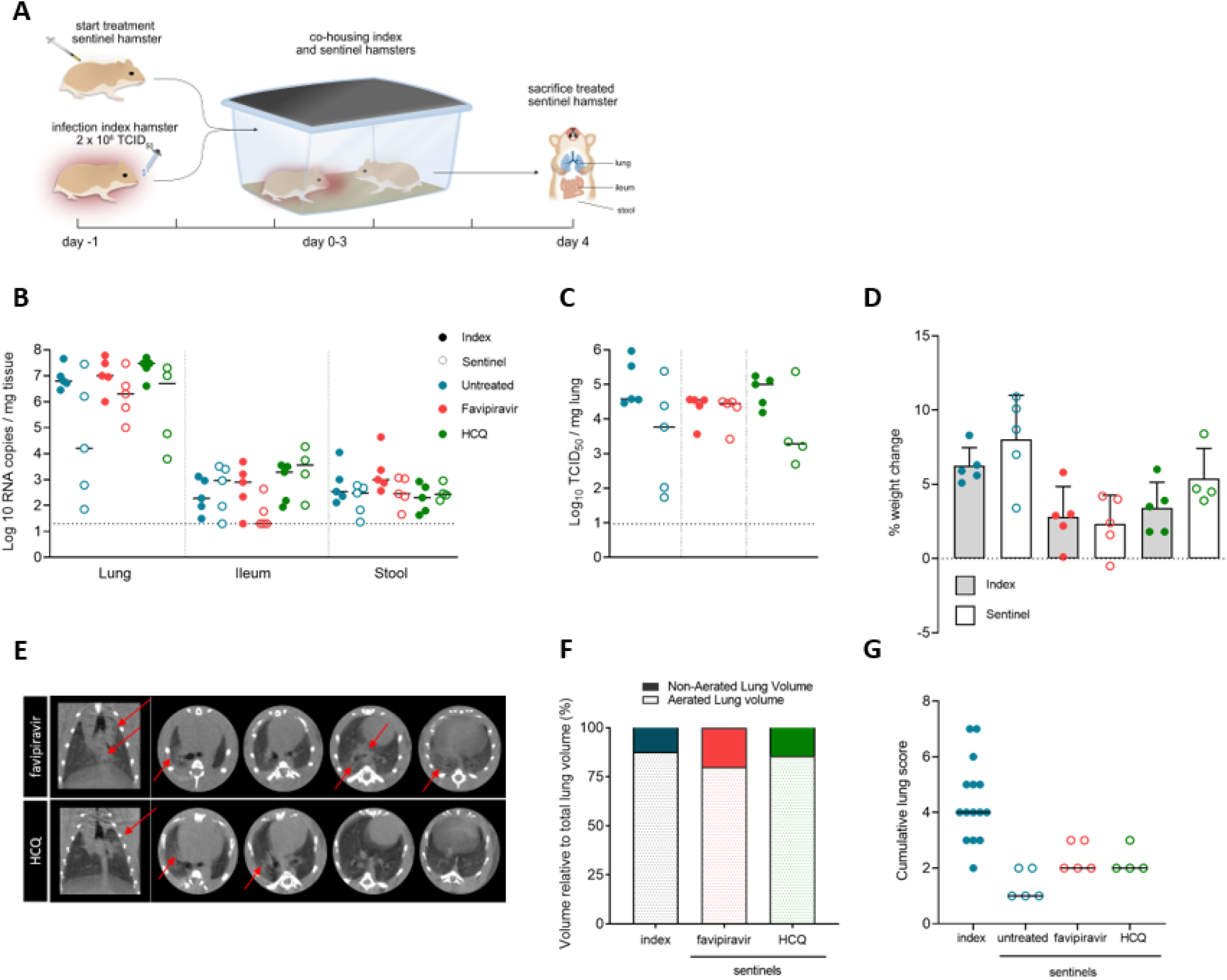
HCQ and favipiravir fail to prevent infection in a direct contact transmission model. **(A)** Set-up of the study. **(B)** Viral RNA levels in the lungs, ileum and stool at day 4 pi are expressed as log_10_ RNA copies per mg tissue. Closed dots represent data from index hamsters (n = 5) inoculated with SARS-CoV-2 one day before co-housing with sentinel animals. Open dots represent data from sentinel hamsters (n = 5 per condition) which were untreated (blue) or treated with either HCQ (green) or favipiravir (red), starting one day before exposure to index animals. **(C)** Infectious viral loads in the lung at day 4 pi/post exposure are expressed as log_10_ TCID_50_ per mg lung tissue. **(D)** Weight change at day 4 pi in percentage, normalized to the body weight at the day of infection (index) or exposure (sentinel). **(E)** Representative coronal and transversal lung μCT images of sentinel favipiravir and hydroxychloroquine (HCQ) treated hamsters at day 4 pi. Red arrows indicate examples of pulmonary infiltrates seen as consolidation of lung parenchyma. **(F)** μCT-derived biomarkers: non-aerated lung volume (reflecting the tissue lesion volume) and aerated lung volume relative to total lung volume of index SARS-CoV-2 infected hamsters and untreated, favipiravir and HCQ treated sentinel hamsters. **(G)** Cumulative severity score from H&E staining of index SARS-CoV-2 infected hamsters and untreated, favipiravir and HCQ treated sentinel hamsters.

### Estimation of HCQ total lung and cytosolic lung concentrations

Based on the measurement of trough concentrations of HCQ at sacrifice (n=14), a mean ± SD trough plasma concentration of 84 + 65 ng/mL (0.251 ± 0.19 μM) was found (Fig 4A). This is comparable to the plasma trough concentrations that were detected in cynomolgus macaques (treated with a dosing regimen of 90 mg/kg on day 1 pi (loading dose) followed by a daily maintenance dose of 45 mg/kg)^14^ and in patients (3-5 days after starting treatment with 200 mg three times daily)^14^. The peak viral load in the lungs was not significantly associated with plasma HCQ concentrations in individual hamsters (Fig 4B), suggesting that a higher HCQ exposure did not result in a more pronounced reduction of viral infection.

**Figure 4.**
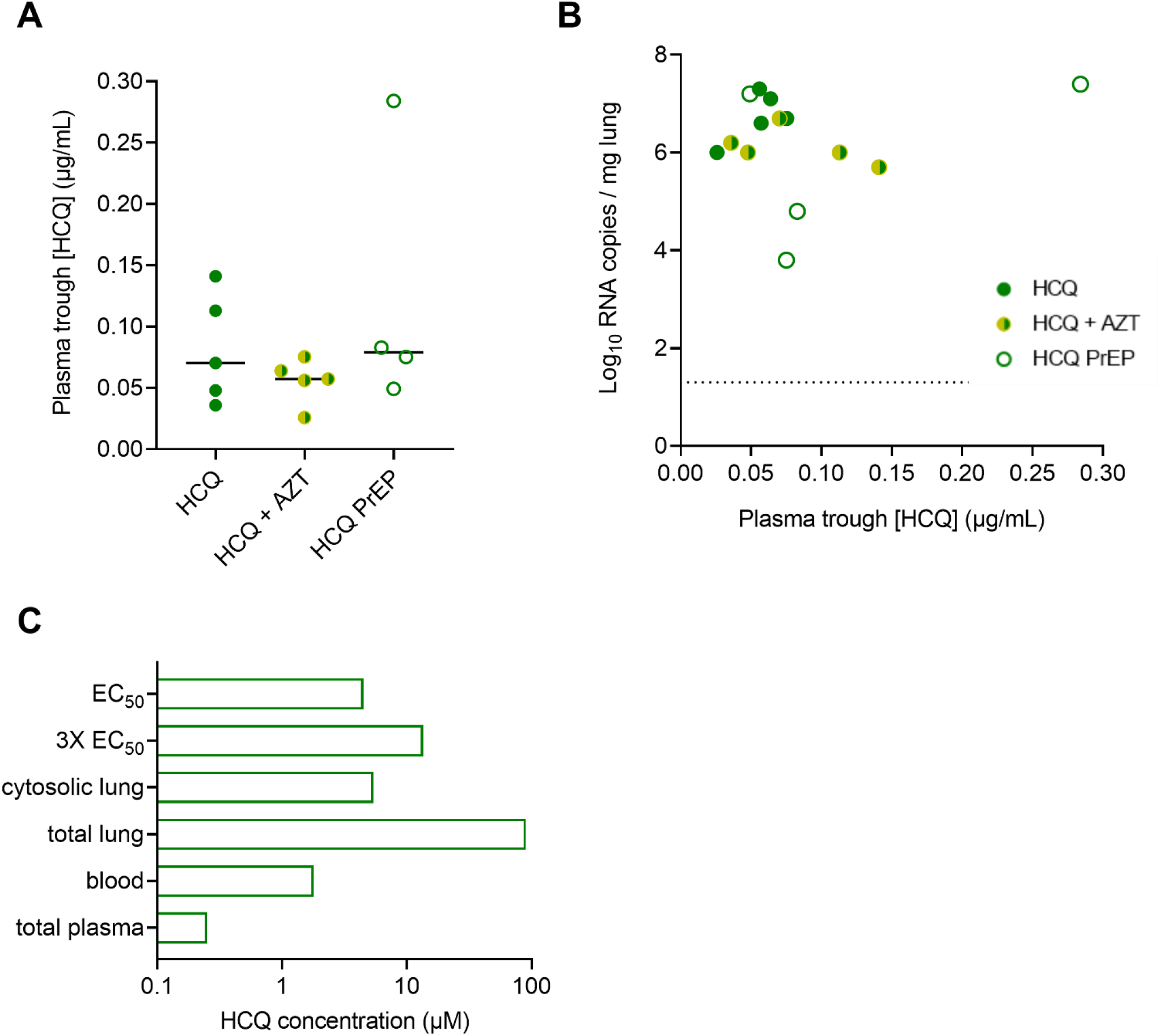
Pharmacokinetics of HCQ in infected and sentinel hamsters. **(A)** Individual plasma trough concentrations of HCQ in hamsters treated with HCQ or HCQ and azithromycin (n=14). **(B)** Viral RNA levels in lung tissue at day 4 pi to HCQ plasma trough concentrations of individual hamsters. **(C)** Summary of trough blood and tissue levels of HCQ in hamsters dosed with 50 mg/kg HCQ sulphate and comparison with *in vitro* EC_50_ values.

According to Equation 1, a whole blood concentration of 1.804 ± 1.39 μM was calculated (Fig 4C). Subsequently, applying Equation 2, this resulted in a total lung concentration of 90.18 ± 69.42 μM, indicating that the lung tissues achieved HCQ concentrations above the reported *in vitro* EC_50_ values, ranging from 0.72 to 17.31 μM, with a median value of 4.51 μM and an interquartile range of 5.44 (25-75%)^29^. To estimate 90% of inhibition of viral replication (EC_90_), the EC_90_ was equated to 3 times the EC_50_, resulting in a target lung concentration of 13.53 ± 16.31 μM. In this case, the efficacy target at trough would be reached when applying this dosing regimen (i.e., 50 mg HCQ sulphate/kg/day). However, it is important to note that the total lung tissue concentrations described above consist of both intracellular and interstitial HCQ concentrations. As the *in vivo* antiviral mechanism(s) of action of HCQ against SARS-CoV-2 has not been clarified yet and might not be exclusively by inhibition of endosome acidification^30^, HCQ concentrations were calculated in cytosolic lung tissue, in the endosomal-lysosomal compartment of cells and in the interstitial compartment. Assuming that cytosolic HCQ concentrations are only 6% of total tissue concentrations, a total cytosolic lung tissue concentration of 5.41 ± 4.17 μM was calculated. This value was in line with the median *in vitro* EC_50_ value, but is well below the estimated EC_90_ value. Also the interstitial concentration was calculated to be 5.41 μM. In contrast, the endosomal/lysosomal HCQ concentration was calculated to be 1.9 mM, which is much higher than the estimated EC_90_.

## Discussion

In a previous study, we showed that wild-type Syrian hamsters are highly susceptible to SARS-CoV-2 infections^16^. Here, we further characterized the hamster infection model to allow the use of this model for antiviral drug evaluation. In agreement with previous studies, upon intranasal inoculation, we observed that the virus replicates efficiently to peak levels (~6 log_10_ TCID_50_/mg) in the lungs on day 4 pi., which is supported by radiological and pathological evidence. Although the virus was also present in the ileum and stool of infected hamsters, levels were significantly lower (~2.5 log_10_ copies/mg). Besides serving as efficient replication reservoirs of SARS-CoV-2, the hamsters also efficiently transmit the virus to co-housed sentinels^15,18^. Here, we demonstrated that the virus is mainly transmitted via direct contact and only to a limited extent via the fecal-oral route. The variability observed in the virus titers in the lungs of the sentinels is probably due to differences in the infection stage of the animals.

Besides hamsters, a variety of other animals have been tested for their permissiveness to SARS-CoV-2, of which ferrets and non-human primates were the most sensitive ones^31–:35^. In ferrets, infectious SARS-CoV-2 was only detected in the nasal turbinate and to a lesser extent in the soft palate and tonsils, but not in the lungs^35^. Although, in a different study infectious virus in the lungs of ferrets was detected, levels remained close to the limit of detection^33^. This indicates that ferrets support SARS-CoV-2 replication, albeit to a lesser extent than hamsters. In SARS-CoV-2-infected macaques (both rhesus and cynomolgus) virus levels were the highest in nasal swabs and the lungs^32,34^. SARS-CoV-2 infection resulted in moderate transient disease in rhesus macaques, whereas cynomolgus macaques remained asymptomatic, but did develop lung pathology as seen in COVID-19^34^. Although aged macaque models may represent the best models for studying more severe COVID-19 disease^36^, both the high costs and ethical considerations (leading to small group sizes) are major drawbacks of non-human primate models. The efficient SARS-CoV-2 replication in the lungs of hamsters combined with development of lung pathology endorses the use of hamsters over any other small animal infection model for preclinical evaluation of the efficacy of antiviral drugs and immune-modulating agents. Potent reduction of SARS-CoV-2 replication in hamsters has been demonstrated by a single dose with a single-domain antibody from a llama immunized with prefusion-stabilized coronavirus spikes^16,37^, thereby validating the use of hamsters to evaluate treatment options against SARS-CoV-2. In addition, our data also indicate that hamsters are highly amenable for studying the potential antiviral effect of small molecules on virus transmissibility in a pre- and post-exposure setting.

In an effort to contribute to the debate on the efficacy of (hydroxy)chloroquine and favipiravir in COVID-19 patients, we evaluated both re-purposed drugs in our hamster infection and transmission model. Treatment with HCQ or combined treatment with azithromycin was not efficacious in significantly lowering viral RNA levels and infectious virus titers in the lungs of SARS-CoV-2-infected hamsters. Lack of efficacy was also demonstrated in the transmission model whereby sentinel hamsters were treated prophylactically prior to exposure to infected hamsters. In SARS-CoV-2 infected ferrets, HCQ treatment was also not able to significantly reduce *in vivo* virus titers^33^. In addition, a recent study in SARS-CoV-2-infected cynomolgus macaques showed that HCQ alone or in combination with azithromycin did not result in a significant decrease in viral loads, both in a therapeutic and in a prophylactic setting^14^. On the other hand, clinical trials with HCQ for the treatment of COVID-19 patients have resulted in conflicting results and controversy. This is especially the case with clinical studies conducted in the early stage of the pandemic, which were mostly small anecdotal studies. Results of large, placebo-controlled, randomized trials are now becoming available. A randomized trial of HCQ as post-exposure prophylaxis after high-to-moderate-risk exposure to COVID-19 showed that high doses of HCQ did not prevent SARS-CoV-2 infection or disease similar to COVID-19^38^. In the RECOVERY trial, a large UK-based clinical study to evaluate potential therapies, HCQ treatment did not result in a beneficial effect in mortality or hospital stay duration in patients hospitalized with COVID-19^39^. These data are in agreement with our results in the hamster model and clearly underline the importance of preclinical studies in animal models in the drug development/re-purposing process.

The lack of effect observed for HCQ in this study and potentially also in other studies may be explained by a pharmacokinetic failure. High lung concentrations of HCQ are caused by massive accumulation (‘ion trapping’) of the compound in acidic lysosomes, which is driven by a pH gradient between cytosol (pH 7.2) and lysosomes (pH 5). However, taking into account the pH partition theory and considering the relative volumes of lung cellular and interstitial compartments, only 6% of total HCQ concentrations in lung tissue is present in the cytosol of lung cells. The other 94% of HCQ is present in the interstitial compartment and intracellularly in lysosomes/endosomes or other subcellular fractions, or bound to proteins. Starting from the measured trough concentrations from treated hamsters at day 4 or 5, the calculated HCQ concentration in the endosomal compartment was 1.9 mM, which would be well above the EC_90_ target. In contrast, cytosolic concentrations in the lung were only slightly higher than the EC_50_ values reported in the literature, and far below the EC_90_ target. Although alkalization of endosomes has been proposed as one of the key mechanisms of the broad-spectrum antiviral effect of HCQ, the mechanism of action against SARS-CoV-2 has not been completely unraveled^30^. Therefore, the very low cytosolic concentrations of HCQ in the lung may explain the absence of an antiviral effect of HCQ against SARS-CoV-2 *in vivo*. Increasing the HCQ dose to reach the EC_90_ might not be feasible in terms of safety, as it may lead to an increased risk of QTc prolongation and fatal arrhythmia. In future studies, lung tissue distribution of (re-purposed) antiviral drugs should be taken into account, along with specification of the subcellular target site, as recommended by Wang and Chen^40^.

In contrast to HCQ, favipiravir was able to inhibit virus replication in intranasally infected hamsters, but the effect was modest and only statistically significant at the viral RNA level. In the transmission model on the other hand, favipiravir failed to reduce viral replication when given as a prophylaxis. This suggests that the antiviral effect of favipiravir in COVID-19 patients will most likely be limited. Also, the efficacy of favipiravir as a pre- or post-exposure prophylaxis seems very modest. Clinical trials to evaluate the potency of favipiravir against SARS-CoV-2 are currently ongoing in China, Italy and the UK^41^. Prior, an open-label, randomized study already showed that in COVID-19 patients with mild symptoms (fever and respiratory symptoms without difficulties in breathing) the clinical recovery rate at day 7 was higher in the favipiravir-treated group compared to the control group, which received treatment with arbidol^42^. However, for COVID-19 patients with hypertension and/or diabetes as well as critically ill patients, the clinical recovery rate was not significantly different between groups, suggesting that favipiravir might be useful for patients with mild symptoms, but not for severely ill patients. One concern with favipiravir is that it has been reported that the trough concentrations (after 8-12h) in critically ill patients are lower than those in healthy persons and do not even reach the *in vitro* obtained EC_50_ value against SARS-CoV-2^43,44^. This unfavorable PK profile of favipiravir has been previously observed in Ebola virus-infected patients^45^. While favipiravir might be well tolerated and safe in a short-term treatment, safety concerns remain as the drug proved to be teratogenic^46^. Therefore, potential widespread use of favipiravir to treat COVID-19 patients should be handled with caution.

In conclusion, we here characterize our hamster infection and transmission model to be a robust model for studying the *in vivo* efficacy of antiviral compounds. Our data endorse the use of Syrian hamsters as the preferred small animal model for preclinical evaluation of treatment options against SARS-CoV-2. Our results also indicate that in both a therapeutic and a prophylaxis scenario, a highly potent antiviral is necessary for a positive outcome. The information we acquired using this model on HCQ and azithromycin is of critical value to those designing (current and) future clinical trials. Of note, in a non-pandemic situation, based on the pre-clinical data we provide, together with the earlier studies in ferrets and non-human primates, there would be no indication to initiate clinical trials with either compound. We recognize the exceptional situation the world is currently in and that clinical trials were initiated at a time when no pre-clinical data was available. However, at this point, the pre-clinical data obtained by us and others on HCQ and azithromycin provide no scientific basis for further studies in humans with these molecules. The very modest reduction of viral load in the lungs of hamsters treated with favipiravir and the lack of efficacy in the transmission model, also suggests that the potential benefit of this drug in humans may be limited as well. Finally, we emphasize the need to develop highly specific, potent and safe pan-corona antiviral drugs. Highly potent drugs are available to treat other viral infections (such as with herpesviruses, HIV, HBV, HCV and influenza virus) and it will without any doubt be possible, given sufficient efforts, to develop also coronavirus inhibitors. Small animal infection models, such as the hamster model, should have a pivotal place in (de)selecting drugs for clinical development.

## Acknowledgments

We thank Kathleen Van den Eynde for excellent technical assistance. We thank Molecubes and Bruker Belgium for their support with the implementation of the micro-CT installation, Jef Arnout and Annelies Sterckx (KU Leuven Faculty of Medicine, Biomedical Sciences Group Management) and Animalia and Biosafety Departments of KU Leuven for facilitating the studies.

This project has received funding from the Covid-19-Fund KU Leuven/UZ Leuven and the COVID-19 call of FWO (G0G4820N), the European Union’s Horizon 2020 research and innovation program under grant agreements No 101003627 (SCORE project), funding from Bill and Melinda Gates Foundation under grant agreement INV-00636, the Stichting Antoine Faes.

G.V.V. acknowledges grant support from KU Leuven Internal Funds (C24/17/061). C.C. was supported by the FWO (FWO 1001719N). S.J. is supported by a PhD fellowship of the Fund for Scientific Research Flanders (FWO). S.t.H. is supported by a KU Leuven internal project. B.H. is a postdoctoral fellow of the Flemish Research Council (FWO - 12R2119N).

