## Supplemental figure 1 for "Antiviral treatment of SARS-CoV-2-infected hamsters reveals a weak effect of favipiravir and a complete lack of effect for hydroxychloroquine"

### Supplemental Information

#### Supplemental Figure 1. SARS-CoV-2 transmission by fecal-oral route is not efficient

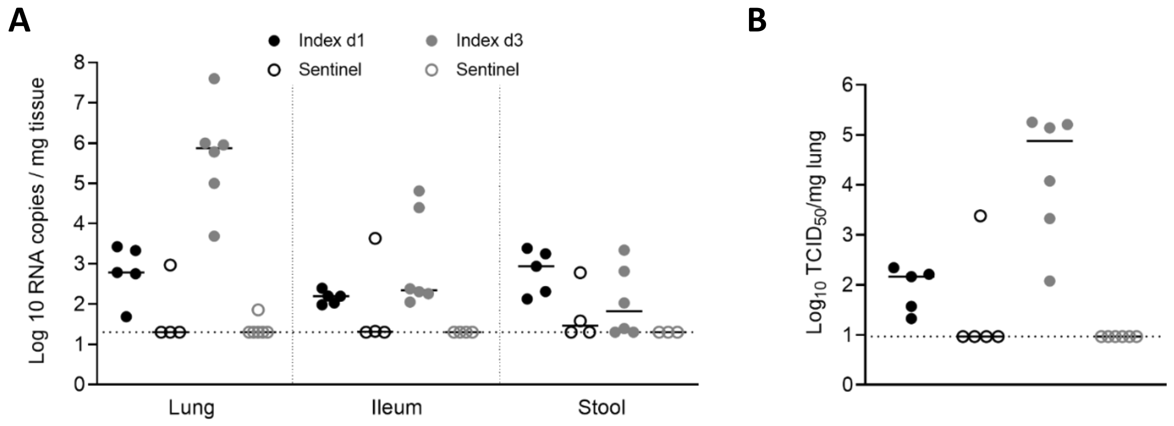


**(A)** Viral RNA levels in the lungs, ileum and stool of index hamsters at day 1 (black) and day 3 pi (grey) and of sentinel hamsters that were exposed for 4 days to the feces of the index hamsters. Viral RNA levels were quantified by RT-qPCR. **(B)** Infectious viral load in the lung of index hamsters at day 1 (black) and day 3 pi (grey) and sentinel hamsters at day 4 post fecal exposure expressed as $TCID_{50}$ per mg of lung tissue.
